# Dysregulation of alternative splicing patterns in the ovaries of reproductively aged mice

**DOI:** 10.1101/2025.05.19.654918

**Authors:** Nehemiah S. Alvarez, Pavla Brachova

## Abstract

Female reproductive aging is characterized by progressive deterioration of ovarian function, yet the molecular mechanisms driving these changes remain incompletely understood. Here, we used long-read direct RNA-sequencing to map transcript isoform changes in mouse ovaries across reproductive age. Comparing young and aged mice after controlled gonadotropin stimulation, we identified widespread alternative splicing changes, including shifts in exon usage, splice site selection, and transcript boundaries. Aged ovaries exhibited increased isoform diversity, favoring distal start and end sites, and a significant rise in exon skipping and intron retention events. Many of these age-biased splicing events altered open reading frames, introduced premature stop codons, or disrupted conserved protein domains. Notably, mitochondrial genes were disproportionately affected. We highlight *Ndufs4*, a mitochondrial Complex I subunit, as a case in which aging promotes the alternative splicing of a truncated isoform lacking the canonical Pfam domain. Structural modeling suggests this splice variant could impair Complex I function, resulting in increased ROS production. Our data suggest a mechanistic link between splicing and mitochondrial dysfunction in the aging ovary. These findings support the model of the splicing-energy-aging axis in ovarian physiology, wherein declining mitochondrial function and adaptive or maladaptive splicing changes are intertwined. Our study reveals that alternative splicing is not merely a byproduct of aging but a dynamic, transcriptome-wide regulatory layer that may influence ovarian longevity. These insights open new avenues for investigating post-transcriptional mechanisms in reproductive aging and underscore the need to consider isoform-level regulation in models of ovarian decline.

**In brief:** Reproductive aging is associated with changes in alternative splicing patterns in the mouse ovary. Our study identifies alterations in exon usage that may have altered protein function and important roles in ovarian physiology as well as in reproductive aging.

## Introduction

### Metabolic challenges in the aging ovary

Female reproductive aging is one of the earliest manifestations of organismal aging, marked by a steady decline in fertility beginning in the mid-30s and culminating in menopause by the early 50s. This decline is driven not only by the depletion of ovarian follicles but also by a pronounced reduction in oocyte quality with advanced maternal age (Bao *et al*., 2024). The molecular mechanisms underlying age-related oocyte dysfunction remain incompletely understood, but accumulating evidence implicates mitochondrial dysfunction and energetic insufficiency in both oocytes and their surrounding somatic cells as key contributors. Mitochondria are critical for oocyte development and survival, providing the ATP needed for processes like meiotic spindle assembly and chromosome segregation. Consistent with this, aged oocytes often exhibit impaired ATP production and elevated oxidative stress, leading to spindle instability and increased chromosomal segregation errors during meiosis (Ju *et al*., 2024). Mitochondrial dysfunction has indeed been identified as a hallmark of ovarian aging: a decline in mitochondrial DNA copy number and respiratory capacity has been correlated with oocyte quality loss and accelerated ovarian cellular senescence (Ju *et al*., 2024). In the follicular niche, the aging microenvironment further exacerbates these issues; senescent granulosa cells in aged ovaries show limited mitochondrial energy output and undergo increased apoptosis, creating an inflammatory, suboptimal milieu for the oocyte (Ju *et al*., 2024). Notably, a recent study of human oocytes from advanced-age women confirmed significant mitochondrial defects and an apparent shift toward alternative energy substrates in these cells (Smits *et al*., 2023). Together, these findings highlight the heavy energetic demands on the ovary and suggest that reproductive aging imposes metabolic stress on ovarian cells. This raises the question of how ovarian cells might adapt at the molecular level to maintain energy homeostasis as mitochondrial function declines with age.

### The splicing-energy-aging axis

Emerging evidence from other systems suggests that one adaptive strategy cells use to cope with metabolic stress is the dynamic regulation of mRNA splicing (Ferrucci *et al*., 2022). In fact, it has been hypothesized that there exists an “energy-splicing-resilience axis” of aging, wherein changes in mitochondrial energy status actively drive changes in pre-mRNA splicing, and this splicing plasticity in turn helps cells preserve energetic homeostasis during aging (Ferrucci *et al*., 2022). Ferrucci and colleagues articulated this concept, proposing that when cellular energy supplies are ample, the splicing program remains in a basal state, but under conditions of mitochondrial impairment and ATP shortfall, cells broaden their repertoire of transcript isoforms as a resilience mechanism (Ferrucci *et al*., 2022). In this model, alternative splicing is upregulated to produce isoforms that can increase energy production or reduce energy expenditure for nonessential processes, partly through signaling cascades that engage energy sensors like AMP-activated protein kinase (AMPK) (Ferrucci *et al*., 2022).

Empirical studies across tissues lend support to this splicing-energy-aging axis. In neuronal cells, for example, inducing mitochondrial damage (and consequent ATP depletion) triggers widespread shifts in splicing patterns, universally biasing transcripts toward shorter isoforms, while constitutively spliced exons remain largely unchanged (Maracchioni *et al*., 2007). Importantly, this splicing modulation in neurons has been shown to correlate specifically with ATP loss rather than with oxidative damage per se (Maracchioni *et al*., 2007), underscoring that the splicing machinery is directly sensitive to cellular energy levels. In skeletal muscle, age-related splicing changes have been tied to the energetic capacity of individuals: older adults with low fitness (indicative of reduced muscle oxidative capacity) exhibit an upregulation of splicing activity and greater alternative isoform diversity compared to high-fitness older adults (Donega *et al*., 2025). The splicing variants enriched in low-fitness muscle were associated with pathways like RNA-protein granule assembly, hinting at a compensatory remodeling of gene expression under energy-poor conditions (Donega *et al*., 2025). The immune system shows a similar linkage between metabolism and splicing regulation. During T-lymphocyte activation, when cellular energy demand sharply rises, there is increased expression and stability of certain splicing regulators (e.g. CELF2) that in turn drive broad alternative splicing changes supporting T cell survival and function (Choi *et al*., 2022). At a gene-specific level, a classic illustration of metabolic-splicing crosstalk is provided by the *PKM* gene: alternative splicing of *PKM* yields the PKM1 and PKM2 protein isoforms, which dictate whether glucose is primarily metabolized through oxidative phosphorylation or diverted to aerobic glycolysis, respectively (Biamonti *et al*., 2018). This well-known switch in pyruvate kinase isoforms demonstrates how splicing decisions can rewire core metabolic pathways to meet cellular energy needs. Furthermore, direct manipulation of energy-sensing pathways can influence splicing outcomes. Activation of AMPK, central energy gauge of a cell, has been shown to modulate splicing factor activity; for instance, in a progeroid aging model, pharmacological AMPK activation (using metformin) inhibited the splicing factor SRSF1 and thereby reduced the production of a toxic alternative splicing product of LMNA (the progerin isoform), alleviating some aging-associated defects (Finley, 2018). Collectively, these examples illustrate that alternative splicing networks respond to energetic and mitochondrial stress across diverse cell types. Rather than being a passive consequence of aging, splicing changes often appear to act as a compensatory mechanism; a means to adjust metabolic gene expression, optimize energy utilization, and maintain homeostasis when cells are under energetic duress (Ferrucci *et al*., 2022).

Given this backdrop, it is plausible that the ovary might also engage a splicing-mediated adaptive program as it ages, in order to cope with mounting mitochondrial dysfunction. However, the contribution of alternative splicing to the ovarian aging process remains unexplored. Recent transcriptome studies of ovarian aging (including single-cell RNA-sequencing efforts) have catalogued changes in gene expression with age, but these analyses have been limited to gene-level measurements and lack information on full-length isoforms due to the short-read sequencing technologies used. As a result, potential age-related shifts in specific transcript isoforms, such as those involving subtle exon usage changes, alternative splicing junctions, or 5′/3′ UTR length remodeling, could not be resolved. To overcome this limitation and directly address the role of alternative splicing in reproductive aging, we employed a direct RNA-sequencing approach using the Oxford Nanopore platform. This long-read, single-molecule sequencing technology allows us to read intact native mRNA molecules, thereby capturing complete transcript isoforms with base-level resolution. In the present study, we applied direct RNA-sequencing to ovaries from young and aged mice to comprehensively characterize age-dependent changes in the ovarian transcriptome at the isoform level. This approach enabled us to identify differential exon and intron inclusion events, shifts in splice site usage, and alternative first/last exon selections (“transcript boundary” remodeling) that emerge with advanced reproductive age. We hypothesized that reproductive aging would be accompanied by specific alternative splicing alterations, potentially as an adaptive response to declining mitochondrial function, and our objective was to map these isoform changes to gain insight into how the ovarian gene expression program is modulated during aging. By delineating the splicing landscape of young vs. aged ovaries in unprecedented detail, our study sets the stage for understanding whether an energy-linked splicing axis is active in the ovary and how such post-transcriptional changes might influence ovarian function and the decline of fertility with age.

## Materials and Methods

### Ovary collection and RNA isolation

Young (6 week-old) C57BL/6 female mice were purchased from Jackson Laboratory, while reproductively aged (14 month-old) C57BL/6 mice were from National Institutes of Aging. 14 month-old mice were selected as the reproductively aged cohort because significant reduction in egg quality and changes in the ovarian microenvironment occur at that age (Chiang *et al*., 2010; Briley *et al*., 2016). Groups of young (3 mice) and aged mice (3 mice) were given an I.P. injection of 5 international units pregnant mare’s serum gonadotropin to synchronously stimulate follicle growth (PMSG, Catalog # 493-10, Lee Biosolutions) 46 hours prior to harvest of whole ovaries. Groups of young (3 mice) and aged mice (3 mice) were given an I.P. injection 5 international units PMSG and then with 5 international units human chorionic gonadotropin (hCG) 46 hours later; ovaries were collected 16 hours later as previously described (Brachova *et al*., 2019). Ovaries were cleaned from bursal sacs and placed in TRIzol (Invitrogen) for RNA isolation according to the manufacturer’s protocol. Mice were maintained in environmentally controlled facilities at the Eastern Virginia Medical School (EVMS) animal facility in a room with a 12-hour light, 12-hour dark cycle (7 am to 7 pm) with ad libitum access to food and water. All animal procedures were performed according to an approved Eastern Virginia Medical School’s Institutional Animal Care and Use Committee protocol.

### Direct RNA-sequencing of young and aged ovaries

Direct RNA-sequencing libraries were prepared for one ovary from each mouse for each age and condition using a Oxford Nanopore Direct RNA-sequencing kit (SQK-RNA002, ONT) the manufacturer’s protocol. Briefly, polyadenylated (poly(A)) RNA was purified from total RNA using a Magnetic mRNA Isolation Kit (NEB, S1550S). 100 ng of poly(A) RNA was annealed and ligated to an oligo-d(T) sequencing adaptor that anchors to the 3′ end of RNA T4 DNA Ligase (NEB, M0202). Adaptor ligated libraries were reverse transcribed using Superscript IV (ThermoFisher, 18090010) and purified with RNAClean XP beads (Beckman, A63987). Twelve libraries were loaded onto individual flow cells (FLO-MIN106D, ONT) and sequenced on a MinION Mk1B (ONT).

### Alignment and isoform quantification

Raw direct-RNA FASTQ files were concatenated to form a single input dataset for uniform processing. The *Mus musculus* reference genome (GRCm38, build 68) and its accompanying Ensembl gene annotation (release 102) were obtained to provide a consistent coordinate framework for read mapping and splice-site correction (Yates *et al*., 2020). Long reads were processed with FLAIR (Full-Length Alternative Isoform Analysis of RNA) (Tang *et al*., 2020). FLAIR’s workflow, comprising *align*, *correct*, *collapse*, and *quantify* stages, first aligns nanopore reads to the genome, refines splice junctions against the annotation, collapses redundant transcripts into non-redundant isoforms, and finally derives transcript-per-million (TPM)–normalized abundance estimates. Stringent splice-checking and annotation-reliant settings were used to maximize isoform confidence, and all operations were parallelized on 120 CPU threads to accommodate the large read volume. The resulting isoform GTF, FASTA, BED, and read-to-isoform mapping files constituted a sample-wide reference transcriptome suitable for downstream analysis.

### In-silico Translation of Isoforms

The collapsed cDNA isoforms were translated *in silico* with bioseq2seq, a transformer-based sequence-to-sequence model trained on mammalian coding sequences (Valencia and Hendrix, 2023). Translations were generated with beam-size 1 to select the single most likely reading frame per transcript; maximum token and decode-length limits prevented over-extension beyond biologically plausible CDS lengths. Predicted peptide sequences were parsed into FASTA format for compatibility with standard annotation tools.

### Protein Domain Annotation

Translated isoforms were scanned against the Protein family (Pfam)-A profile HMM collection (release 35.0) using *hmmsearch* to assign conserved domain annotations and to facilitate functional interpretation of novel splice variants (Mistry *et al*., 2021).

### Preprocessing and Isoform Proportions

To identify genome-wide splicing changes between young and aged ovaries, we implemented a Bayesian hierarchical model to compare isoform usage proportions across conditions. We first grouped raw transcript counts by their promoter of origin and filtered out any promoter group lacking expression in one of the conditions (each group was required to have at least one non-zero count in both young and aged samples). Within each remaining promoter group, transcript counts for each sample were converted to proportions by dividing by the group total count, yielding the fraction of transcripts originating from each isoform. These isoform proportions (analogous to percent-spliced-in values) were bounded away from 0 and 1 (rescaled to lie between 0.001 and 0.999) to avoid numerical issues. This preprocessing ensures that differential splicing is assessed on a relative usage scale, effectively controlling for overall gene expression differences. Notably, using relative proportions addresses the compositional nature of isoform data—changes in an isoform fraction reflect true shifts in splicing, independent of gene-level expression changes.

Classical RNA-seq analysis methods (e.g., edgeR or DESeq2) model absolute read counts with negative binomial distributions and are designed for detecting differential total expression per gene or transcript (Robinson *et al*., 2010; Love *et al*., 2014). These count-based tools are not optimized for compositional data like isoform ratios, as they assume independent features and do not account for the constraint that isoform proportions sum to one within a gene. In fact, applying gene-level methods directly to isoform counts can be problematic: it fails to capture the varying estimation uncertainty across isoforms and may lead to reduced power or inflated false positives (Leng *et al*., 2013). Leng et al. (2013) specifically cautioned that using standard count-based techniques on isoform-level estimates is *“not recommended”* due to these issues, advocating for hierarchical modeling to properly share information between isoforms. Therefore, a Bayesian approach was chosen to naturally accommodate the relative nature of isoform usage data and to better model the inherent uncertainties in isoform proportion estimates.

### Bayesian Model for Differential Isoform Usage

We applied a custom Bayesian Beta-binomial model to assess shifts in isoform ratios between conditions. In our hierarchical model, the proportion values for each isoform (across all replicates and conditions) are treated as random draws from an underlying Beta distribution with isoform-specific shape parameters (α and β). This choice is well-suited for data bounded between 0 and 1, and it flexibly captures different usage biases: for example, an isoform with α >> β has a propensity for high usage (fraction near 1), whereas β >> α would imply the isoform is usually lowly used (fraction near 0). We placed weakly informative Gamma priors on each isoform α and β (shape=5, scale=1), reflecting a mild prior belief that extreme proportions are unlikely without substantial evidence. These priors act as regularization, shrinking estimates toward modest usage levels in the absence of strong data support, which is particularly advantageous given our limited sample size. Such prior-informed smoothing is a key benefit of the Bayesian framework, it helps stabilize inference with small *n* by “borrowing strength” across isoforms and samples (Leng *et al*., 2013). Indeed, empirical Bayes and fully Bayesian methods have demonstrated improved performance in RNA-seq analyses by sharing information in this way (Leng *et al*., 2013; Tiberi and Robinson, 2020).

Posterior sampling was performed using Markov Chain Monte Carlo and yields a posterior distribution for each isoform usage parameters and, by extension, for each isoform difference in usage between conditions. From the posterior samples, we directly computed the log₂ fold-change in isoform proportion (aged vs. young) along with its 95% highest posterior density (HPD) interval. Importantly, the Bayesian approach provides a probabilistic assessment of differential splicing: for each isoform, we calculated the posterior probability that its usage is higher in aged ovaries (and conversely in young ovaries). This probability metric offers an intuitive measure of confidence in the direction of splicing change, as opposed to relying only on p-values or FDR thresholds. We designated an isoform as significantly differentially spliced if two criteria were met: (1) the 95% HPD interval of its log₂ fold-change in proportion did not include zero (i.e. no overlap with “no change”), and (2) the posterior probability of a usage increase in one condition exceeded 0.95. In other words, we require high-confidence that the isoform usage shifted substantially between young and aged samples. This Bayesian decision criterion is analogous to using a stringent significance cutoff, but it has the advantage of incorporating the full posterior uncertainty of the estimate. Methods like MISO and MATS have similarly leveraged Bayesian posterior distributions to call differential isoform usage with high confidence (Katz *et al*., 2010; Shen *et al*., 2012), and more recent Bayesian frameworks have shown that such probability-based thresholds control false discoveries well while improving detection power in practice (Tiberi and Robinson, 2020).

#### Advantages of a Bayesian modeling approach

In summary, adopting a Bayesian hierarchical model for isoform usage offers several key advantages over classical approaches. First, it models proportions directly (using Beta/Beta-binomial distributions) and thus properly accounts for the compositional structure of transcript data, unlike methods built for independent counts. Second, the Bayesian framework accounts for uncertainty in a principled way: rather than outputting only point estimates and p-values, it yields posterior distributions and credible intervals for isoform usage changes, enabling statements like “there is a 98% posterior probability that isoform X usage is higher in aged ovaries.” This quantification of uncertainty is crucial when read mapping ambiguity or limited read depth can obscure isoform quantification (Katz *et al*., 2010; Tiberi and Robinson, 2020). Third, Bayesian modeling (especially with informative priors and hierarchical sharing) often improves performance with small sample sizes. By incorporating prior knowledge and pooling information across genes, it mitigates the instability that plagues frequentist estimates in low-replicate scenarios (Shen *et al*., 2012). Studies have shown that Bayesian or empirical Bayes methods can achieve greater sensitivity and specificity in detecting differential isoform expression, even with few replicates, compared to traditional count-based tests (Leng *et al*., 2013; Tiberi and Robinson, 2020). These advantages motivated our choice of a Bayesian model for differential splicing analysis, as it provides a robust, uncertainty-aware inference of isoform regulation changes between young and aged ovarian samples.

### Alternative Splicing Event Annotation Pipeline

#### Input Transcript Models

The starting point was a collapsed transcriptome GTF produced by FLAIR, containing multi-exon and single-exon transcript models for each gene. All transcripts were grouped by gene, and a custom Python module was used to compare transcript structures and integrate coverage data to identify various classes of alternative splicing events (implemented as custom code). The analysis proceeded gene-by-gene in a strand-aware manner, leveraging both direct GTF structural comparisons and an inferred expression-support signal derived from normalized exon coverage. The following event categories were annotated: alternative promoters, alternative 5′ splice sites, alternative 3′ splice sites, alternative exons (including cassette and mutually exclusive exons), retained introns, and alternative transcription end sites.

#### Alternative Promoters (TSS Clusters)

To identify *alternative promoter* events, transcripts of each gene were first clustered by their transcription start site (TSS). Transcripts with TSS positions within a small genomic window (±1 bp) were considered to share the same promoter cluster. If a transcript TSS did not overlap any existing cluster, it initiated a new promoter group. This ensured that transcripts initiating from distinct genomic positions were assigned to separate promoter groups, representing distinct promoter usage. Each unique TSS cluster was recorded as an alternative promoter for the gene. For genes with multiple promoter clusters, a fractional distance metric was computed to indicate the relative position of each promoter along the gene span (0.0 for the most upstream TSS, up to 1.0 for the most downstream). All transcripts were annotated with their promoter group membership, and those originating from downstream clusters (when multiple clusters exist) were thus classified as alternative promoters.

#### Normalized Exon Coverage Profile

For each gene (and for each promoter-based subgroup of its transcripts), we generated a strand-specific normalized coverage profile across the gene exonic span to help delineate exon and intron features. This profile was derived by superimposing all exons from the transcripts in the group onto a single coordinate axis from the first exon start to the last exon end. A coverage value was calculated at each nucleotide position as the sum of 1/(number of transcripts spanning that exon) for each overlapping exon. In effect, bases that are exonic in all transcripts of the group received a normalized coverage of ∼1.0, whereas bases present in only a subset of transcripts had fractional coverage (<1.0), and intronic regions (absent from all transcript exons) had coverage ∼0. This normalized coverage approach provides an approximate “exon inclusion” frequency across the transcripts. We then applied a peak-finding algorithm to this profile to locate local peaks (high points corresponding to putative exon centers) and valleys (low points corresponding to intronic regions). Peaks represented candidate exonic segments, with their height reflecting the fraction of transcripts including that segment, while valleys indicated boundaries between exons (intron centers with minimal coverage). Each valley position was expanded leftward and rightward to determine the span of the intron (valley region) between flanking exons. This provided approximate intron coordinates for further analysis. The normalized coverage profile, along with the identified peak and valley positions, was then used to support the detection of alternative splicing events as described below.

#### Alternative 5′ Splice Sites (Donor Site Variation)

We defined *alternative 5′ splice site* events (alternate donor sites) as cases where an intron 5′ end occurs at multiple positions across transcripts, resulting in alternative starts of the downstream exon. Using the normalized coverage profile and identified intronic regions, we scanned for abrupt drops in coverage at exon boundaries that would indicate such donor site variation. On the positive strand, the algorithm iterated through exon regions in the 5′→3′ direction and flagged a position *i* as a candidate alternative 5′ site if the coverage at *i* was lower than at the preceding base *i*−1, signaling a step down in coverage within what should be an exon. Importantly, any candidate position overlapping an intronic region or lying beyond the last exon end was ignored. This logic detects a point where one transcript exon ends (entering the intron), while another transcript exon continues further, a hallmark of an alternative donor splice site. For negative-strand genes, the scan was inverted (iterating 3′→5′) to detect drops in coverage when moving in the opposite orientation (since a 5′ splice site is at the genomic downstream end of an intron on the minus strand). By comparing coverage differences across exon–intron junctions in a strand-aware fashion, we pinpointed bases that serve as alternate donor splice junctions supported by the transcript models and coverage dips. We additionally ensured that such events were not misattributed to transcript termination differences by excluding any donor-site candidate that coincided with an alternative transcript end position (see below). Each confirmed position was annotated as an alternative 5′ splice site, and any exon ending at that genomic coordinate was labeled accordingly in the output GTF (feature type exon_alt5).

#### Alternative 3′ Splice Sites (Acceptor Site Variation)

Similarly, *alternative 3′ splice site* events (alternate acceptor sites) were identified as positions where an intron 3′ end (the acceptor junction for the upstream exon) varies between transcripts. For plus-strand transcripts, we examined each coverage peak (exon) and scanned backwards (toward the 5′ direction) from the exon center toward its start, looking for a position where coverage increases from one base to the next (when moving right-to-left). A position j was flagged as an alternative acceptor if coverage at j was lower than at j+1, indicating the point at which an incoming intron coverage rises into the exon – i.e. a transcript that has a longer intron (skipping some exon portion) vs. another that starts the exon at a downstream acceptor j+1. The algorithm skipped over any positions lying inside intronic regions (ensuring the candidate is at an exon boundary) and stopped once it reached another peak or the transcript start. For minus-strand genes, the procedure was symmetric but in the forward direction (scanning from exon center toward the 3′ end) due to reversed transcription orientation. This effectively finds alternate acceptor junctions where one transcript includes extra sequence at an exon 5′ end that another transcript splices out. To avoid conflating alternative 3′ splice sites with alternative TSS events, any candidate acceptor site that coincided with a known alternative promoter (TSS cluster) position was not counted as an alternative 3′ splice site. Verified alternative acceptor positions were annotated in the GTF, and exons beginning at those coordinates were labeled exon_alt3 in the feature field.

#### Alternative Exons (Cassette and Mutually Exclusive Exons)

We annotated *alternative exons* as internal exonic segments that are included in some transcript isoforms but skipped (omitted) in others. This category covers simple cassette exons (one exon optional vs. skipped) as well as mutually exclusive exons (where two alternative exons occupy the same relative position, each present in complementary subsets of transcripts). Detection of alternative exons relied on the normalized coverage profile peak analysis. Specifically, any internal exon that did not appear in all transcripts of the group exhibited a reduced normalized coverage peak (<1.0). We extracted all coverage peak positions and identified those with height significantly below 1.0 (indicating partial exon inclusion). To focus on well-defined exon-skipping events, we further required that such a sub-maximum peak be flanked by pronounced valleys on both sides (i.e. it is an isolated internal exon bounded by introns). In practice, for each peak with fractional coverage, we found the nearest coverage minima (valleys) before and after the peak and retrieved the exact exon coordinates falling between those valley positions. We excluded the first and last exons from this analysis (to distinguish alternative internal exons from alternative promoters or ends). Any exon that met these criteria, an intermediate exon used by only a subset of transcripts, was labeled as an alternative exon. In the updated GTF, such exons were marked with feature type exon_ce (denoting cassette/alternative exon). Notably, mutually exclusive exon pairs would both appear as alternative exons by this method: each alternative exon of the pair shows incomplete inclusion across transcripts (each is missing in the transcripts that use the alternate exon), yielding sub-maximal coverage peaks for both. Thus, the pipeline can capture mutually exclusive exons by labeling each involved exon as exon_ce (if they satisfy the coverage and structural criteria). Each transcript that contains an alternative exon was flagged in its attributes (see Output, below), and transcripts skipping the exon simply lack that exon_ce segment.

#### Retained Introns

Retained intron events were detected by identifying intronic regions that show evidence of being transcribed (not spliced out) in at least one isoform. Two complementary approaches were used. Structural comparison: for each intron (interval between exons) defined by one transcript, the pipeline checked if any other transcript in the gene has a continuous exon that spans across that intron coordinates. If so, this suggests that the region is not spliced out in that transcript, i.e. an intron retention event. Coverage analysis: we leveraged the normalized coverage valleys corresponding to introns. After defining all intron intervals from the coverage minima (valley spans) as described above, we calculated the average normalized coverage within each intron interval. Introns that had non-zero (and especially significant fractional) coverage indicated that some transcripts contribute exonic reads in those normally intronic positions. In other words, if the coverage profile does not drop to zero in an intron, it implies one or more transcripts include that sequence. Such intronic regions were flagged as retained intron candidates. In the output GTF, any exon overlapping a flagged intron region was relabeled with exon_ir to denote that it represents a retained intron sequence. This labeling was achieved by marking all base positions within each identified intron interval and intersecting these with exon coordinates. As a result, an exon that fully or partially covers a normally intronic span is annotated as exon_ir. Each transcript containing one or more retained introns was also annotated in its attributes (incrementing an intron retention count; see below). Conventional introns (with zero coverage and not spanned by any other transcript exon) were not labeled as retained.

#### Alternative Transcription End Sites (Alternative TES)

We identified *alternative transcription end site* events by clustering transcript 3′ end positions (transcription end sites, TES) on a per-gene basis (accounting for strand orientation). For each transcript, the TES was taken as the coordinate of the transcript last exon end if on the + strand, or the first exon start if on the – strand. Transcripts with end sites within a small window (±1 nt) were grouped, meaning their 3′ ends were considered the same cluster. Distinct clusters thus represent different polyadenylation sites or termination positions used by different isoforms. If a transcript end did not fall in the vicinity of any other, it initiated a new cluster. This method effectively merges transcripts ending at essentially the same nucleotide into one group, separating them from transcripts that end elsewhere. Each unique end cluster was recorded as an alternative end (alternative poly(A) site) for the gene. To quantify relative positions of multiple end clusters, a fractional distance along the gene was computed: for plus-strand genes, end sites closer to the gene 3′ end were assigned higher fractional values (approaching 1.0), whereas for minus-strand genes the calculation was inverted (since a smaller genomic coordinate actually lies downstream in transcription). This provided a normalized measure of how far downstream each alternative TES was. Alternative end events were annotated by marking the corresponding transcript end exons in the GTF as exon_alt_end. In practice, the last exon of transcripts using a downstream poly(A) site was labeled exon_alt_end (and similarly for the first exon of a minus-strand transcript using an alternative downstream end). Transcripts were also annotated with their respective end cluster (and fractional position) in the output attributes.

#### Output GTF Annotation and Transcript Attributes

The pipeline outputs an updated GTF file in which exons are relabeled according to the alternative splicing features identified. Each exon feature field (column 3 of the GTF) is replaced with a composite label indicating any alternative splicing event that overlaps that exon coordinates. For example, an exon that was determined to use an alternative 5′ donor site would be labeled exon_alt5, and one with both an alternative acceptor and being an underutilized cassette exon would be labeled exon_alt3_ce (the labels are concatenated with an underscore, all prefixed by “exon”). The possible exon labels include: exon_alt_pro (exon at an alternative promoter TSS, i.e. first exon of a downstream promoter group), exon_alt5 (exon with an alternate 5′ splice site), exon_alt3 (exon with an alternate 3′ splice site), exon_ce (cassette or alt exon), exon_ir (exon representing a retained intron sequence), and exon_alt_end (exon at an alternative transcript end site). These labels were assigned based on the positional overlap of the exon with the features detected (the set of labels for each base position of the exon was aggregated). In addition to relabeled exons, the method appends informative attributes to each transcript entry in the GTF. Each transcript is annotated with a promoter group identifier (indicating which alternative promoter cluster it belongs to) and with numeric attributes reflecting that transcript involvement in various splicing events. For example, the number of alternative donor sites (a5) utilized by the transcript, the number of alternative acceptors (a3), the count of alternative exons it includes (ce), and any retained introns (i), along with the total possible events of each type identified for the gene or promoter group (a5t, a3t, cet, it, etc.), were recorded in the attributes. Transcripts arising from multiple promoter groups have a fractional promoter usage value (ap) and those with multiple end site options have a fractional TES value (ae), corresponding to the relative positions described above. These transcript-level attributes provide a summary of each isoform splicing pattern and its grouping by promoter. The custom code implementing this pipeline writes out the annotated GTF with the modified exon feature labels and transcript attributes (promoter group and splicing event metrics) as the final output (Custom Python Module). All analyses were performed with this in-house Python script, and the methodology described here reflects the exact logic in that code. (No transcript abundance quantification or TPM filtering steps were applied in this annotation pipeline, focusing purely on structural and coverage-based splicing evidence.)

#### Alternative splicing event calling (transcript-centric model)

After isoform collapsing, we treated each full-length transcript, rather than individual splice junctions, as the primary unit for splicing analysis. The workflow proceeds gene-by-gene and relies only on the GTF structure plus an internally derived, strand-aware exon– coverage profile; no junction counts or short-read PSI estimates are used. For every transcript we stored a vector of binary flags indicating which events it includes; the complementary transcripts implicitly exclude those same events.

#### Percent spliced (PS) assignment

Within each promoter group, transcript TPMs were converted to isoform-usage proportions (TPM of a transcript divided by the sum of TPMs of all transcripts that share its promoter). The log_2_ fold change of this proportion between the two biological conditions was taken as that transcript percent spliced (*PS*) value. Because every alternative splicing event is physically embedded in the full-length isoform, the *PS* value of an event is inherited directly from the *PS* of the transcript(s) that include it (and conversely from the excluding transcript for the “skipped” form). Thus each event receives a single, internally consistent magnitude that exactly matches the change in usage of the complete transcript that embodies it.

#### Why a transcript-centric model is preferable for Nanopore direct-RNA data

*Full combinatorial context*. Nanopore reads span the entire mRNA, so alternative promoters, splice sites, alternative exons, intron retention, and poly(A) choices are observed simultaneously in each molecule. Scoring events through whole-isoform proportions preserves these naturally co-occurring combinations and avoids double counting overlapping junctions. *Coherence between splicing and expression*. The *PS* metric is exactly the differential usage of a biologically intact transcript; downstream functional consequences (protein isoforms, UTR changes) are therefore interpreted on the same quantitative scale. *Elimination of incompatible read allocations*. Short-read PSI methods must partition ambiguous junction reads among multiple events, often producing inconsistent or inflated estimates. Long-read isoform proportions bypass this ambiguity entirely. *Promoter-aware normalization*. By computing proportions within promoter groups, variation caused by alternative TSSs is separated from downstream splicing changes, preventing dilution or exaggeration of splice-site effects. *Direct compatibility with coverage-based QC*. The same normalized coverage profile used to discover events also confirms their quantitative support, ensuring that low-depth artefacts are not inflated into false positives.

Together, these features make the transcript-centric, proportion-based *PS* framework the most logical and internally consistent way to quantify alternative splicing with Nanopore direct RNA-sequencing. All event calls and *PS* assignments were produced by the custom Python module available in our public repository.

### Statistical Tests

Figure 2B-E, utilized two-sided exact binomial test for imbalance between up- and down-regulated counts. For Figure 2F–G, distal-vs. proximal-boundary counts in up- and down-regulated (and unaffected) isoforms were compared with two-sided Fisher’s exact tests (2 × 2 tables, Bonferroni-corrected for multiple comparisons). Figure 3 analyses used Pearson’s chi-square tests to compare the proportion of single-vs. multi-isoform genes and two-sided Fisher’s exact tests to assess enrichment of alternative splicing event categories. For Figure 5A and B, chi-squared tests were used to evaluate differences in the distribution of splicing event types contributing to Pfam domain gains or losses between young and aged ovaries after PMSG and PMSG + hCG stimulation.

**Figure 1.**
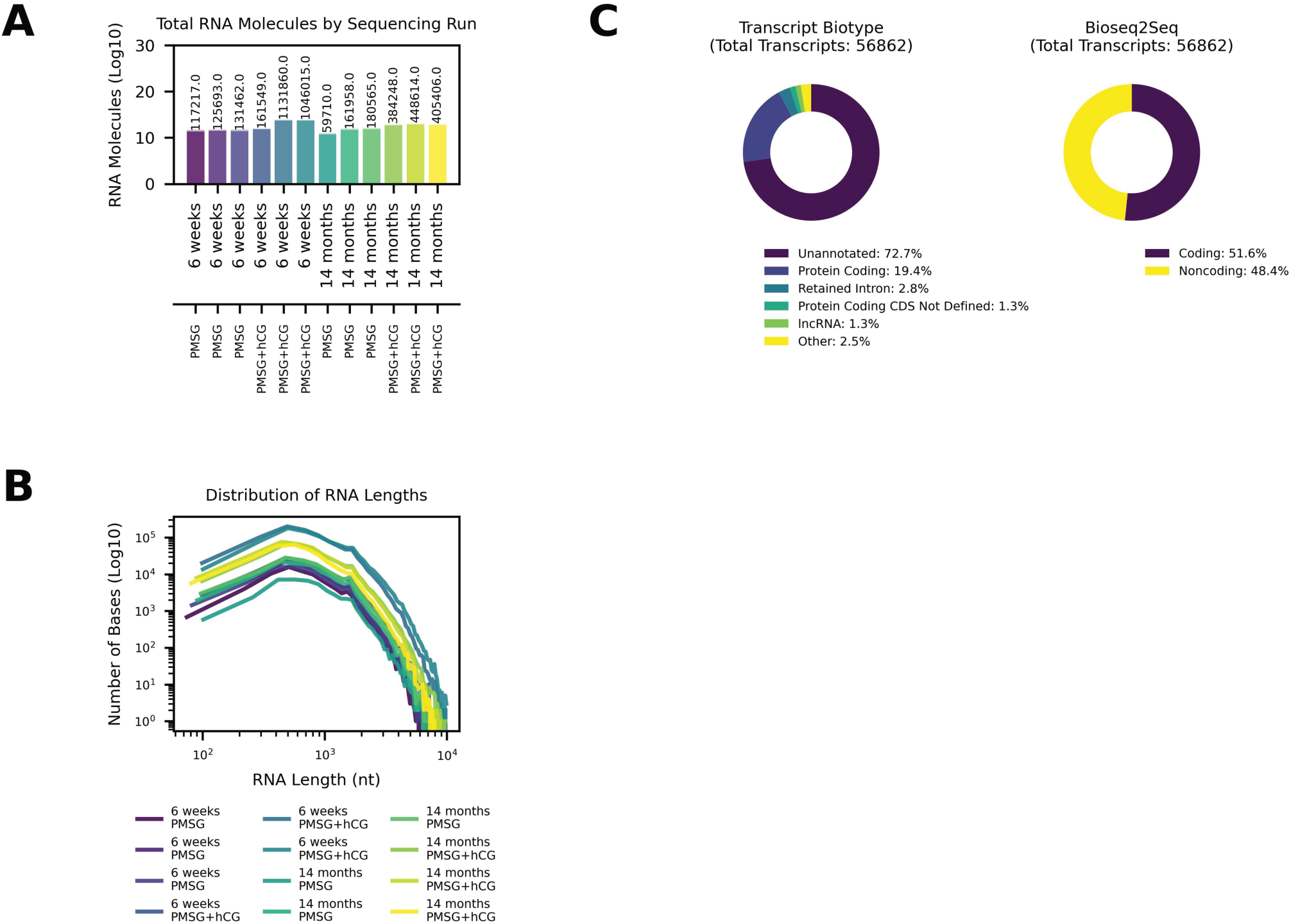
Direct RNA-sequencing yields a high transcript count and reveals novel transcripts in the mouse ovary. (A) Bar plot showing the total number of RNA molecules sequenced in each of 12 runs (y-axis on a log_10_ scale). Bars are colored for ease of viewing, age and gonadotropin stimulation marked on x-axis, and each bar represents an individual sequencing run. (B) Length distribution of sequenced RNA molecules. Each curve corresponds to one experimental condition (6-week or 14-month, + PMSG/ + PMSG + hCG, color-coded as in the figure legend) and plots the total number of bases sequenced (y-axis, log_10_ scale) across transcript lengths (x-axis, in nucleotides). (C) Ovarian transcript composition and coding potential. Left: Donut chart illustrating the proportion of transcripts by Ensembl biotype among all unique transcripts identified. Right: Donut chart classifying these transcripts by predicted coding potential (Bioseq2seq model).

**Figure 2.**
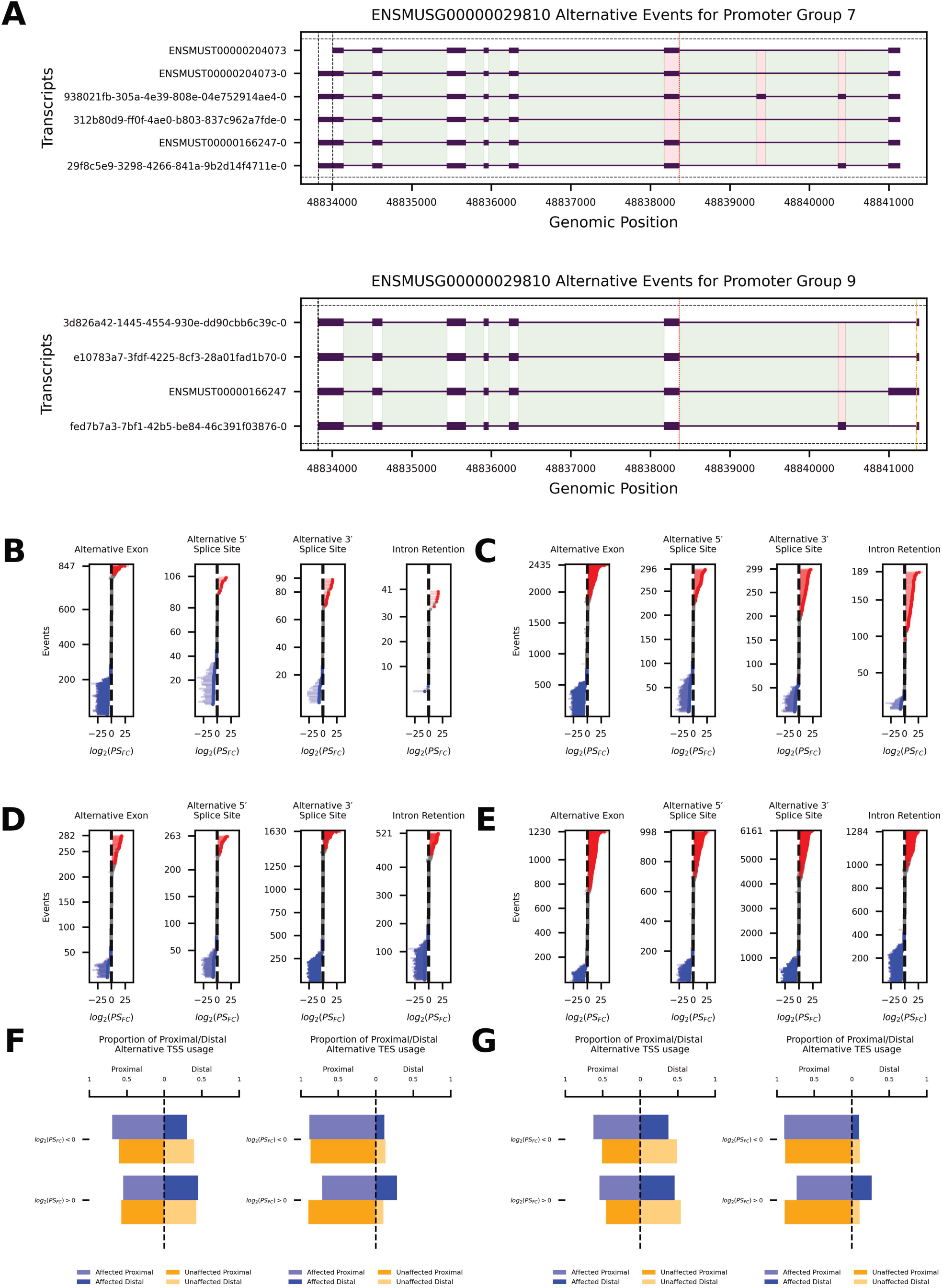
Reproductive aging skews alternative splicing outcomes and transcript boundary selection after gonadotropin stimulation. (A) A promoter group (see Methods) plot for an example gene (Tmem176b; ENSMUSG00000025955) with transcripts from two different promoter groups. Alternative 5′ splice sites are indicated by orange dashed lines, alternative 3′ splice sites by red dotted lines, and alternative transcript end sites by black dash-dot lines. Alternative exons are shown as semi-transparent red rectangles, introns are highlighted in green, and retained introns are marked in dark green. (B–E) Summary of age-associated splicing events (alternative exons, alternative 5′/3′ splice sites, intron retention) for young vs. aged. Panels B and D show inclusion and exclusion events for 6-week vs. 14-month PMSG. Panels C and D show inclusion and exclusion events for 6-week vs. 14-month PMSG + hCG. Y-axis are individual alternative splicing events, x-axis is log_2_ fold change of percent spliced (*PS*). Horizontal red and blue lines are significant events, horizontal grey lines are not significant. (F, G) Proportional usage of proximal vs. distal transcript boundaries significantly affected (blue) or unaffected (orange) isoforms in 6 weeks vs. 14 months after PMSG (F) or PMSG + hCG (G). F and G left sub-panel compares transcription start sites (TSS) and the right sub-panel compares transcription end sites (TES).

**Figure 3.**
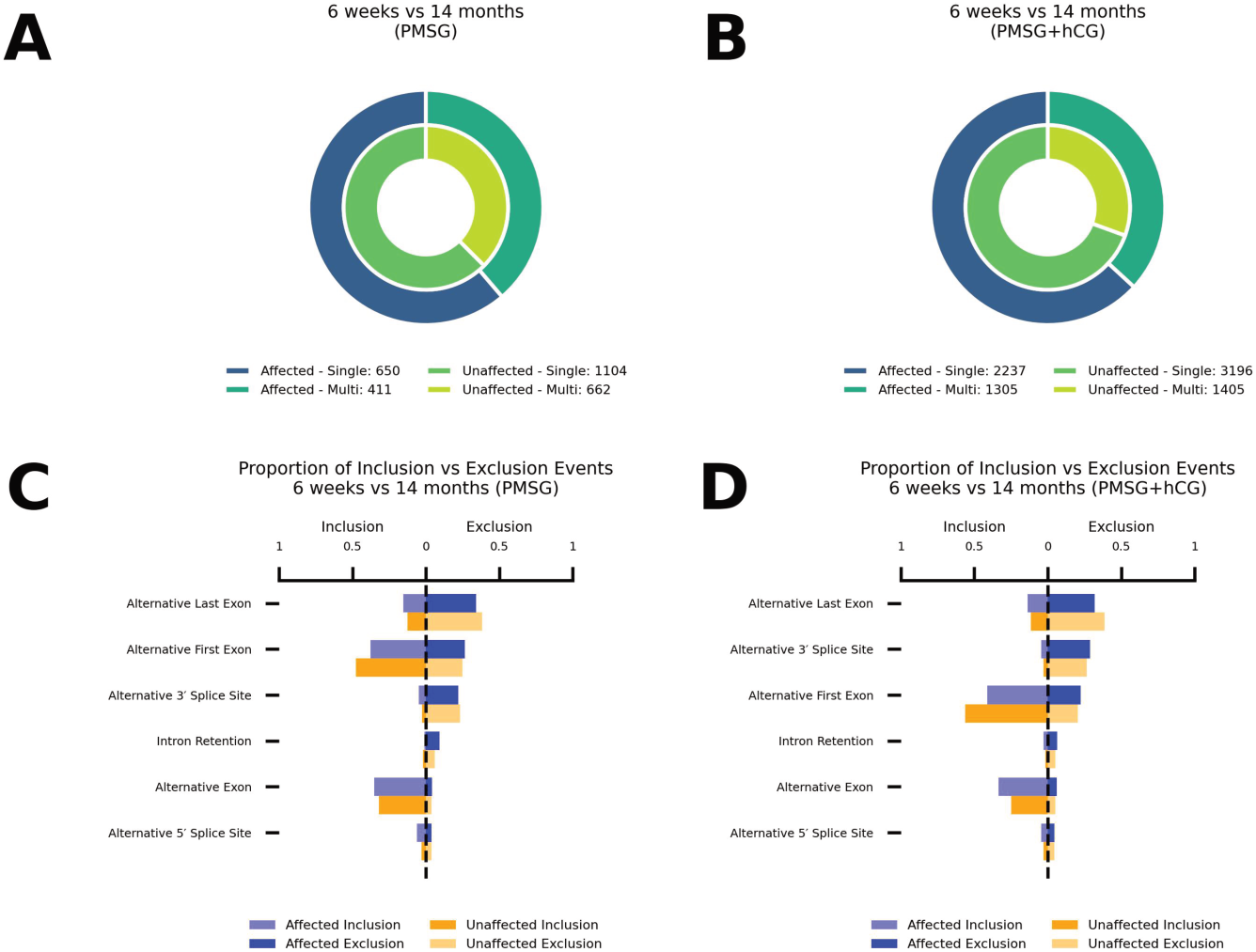
Reproductive aging increases transcript complexity and differentially affects splicing event types. Gene transcript complexity in 6-week vs. 14-month ovaries after PMSG (A) and PMSG + hCG (B). Donut charts show the proportion of single-transcript vs. multi-transcript promoter groups. (C, D) Global distribution of age-biased (“affected”) and age-neutral (“unaffected”) alternative splicing categories. Bar graphs illustrate the proportion of alternative splicing categories that are affected or unaffected in the PMSG (C) and PMSG + hCG (D) conditions. For each panel, the left side represents events resulting in inclusion, and the right side represents events leading to exclusion.

**Figure 4.**
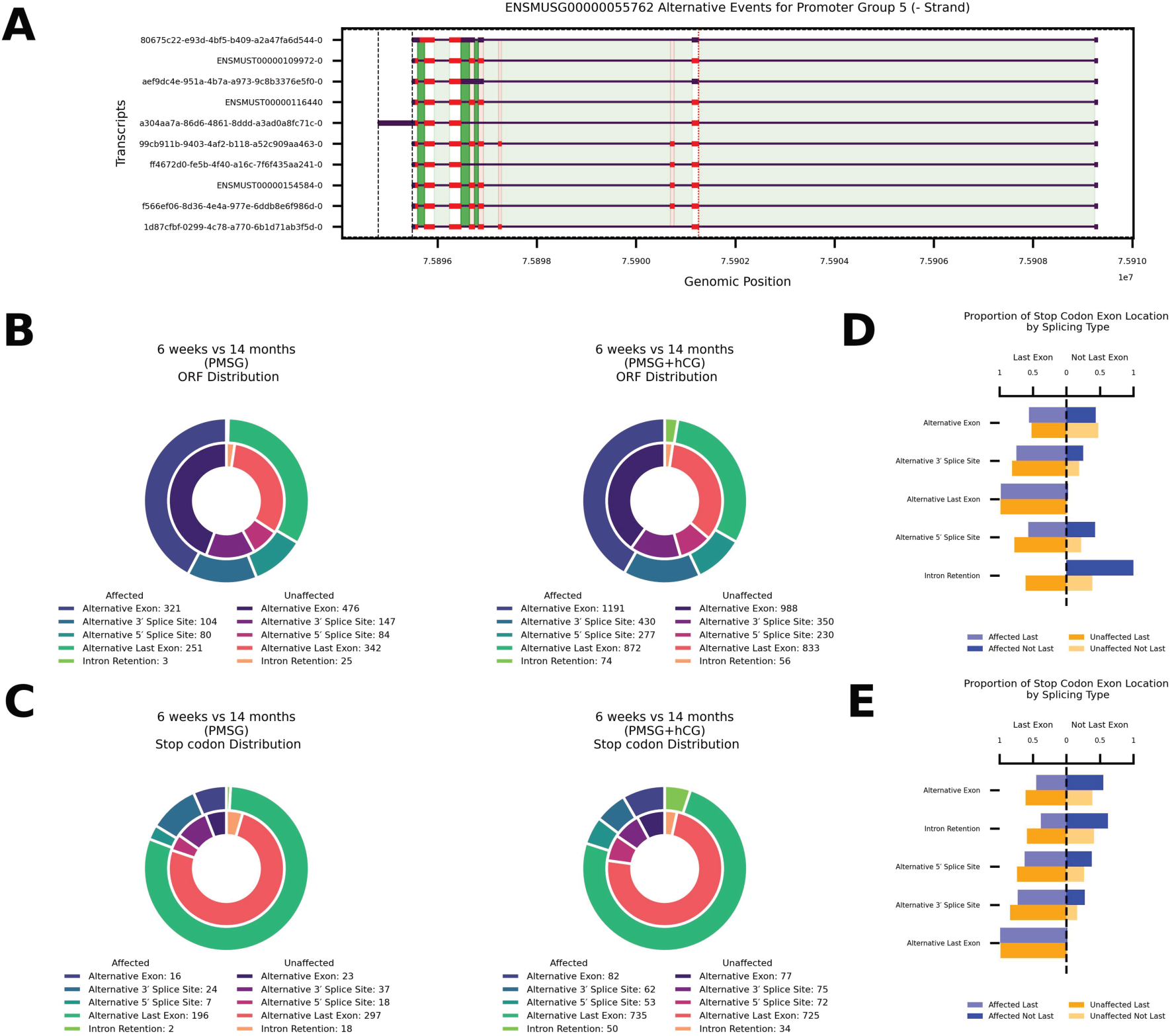
Age-associated changes in alternative splicing alter the protein-coding capacity of the ovary. (A) A promoter group (see Methods) plot for an example gene (*Eef1d*, ENSMUSG00000055762) that illustrates how alternative splicing affects ORFs. Exons are depicted as black boxes and the ORF is shown in red. Alternative 5′ splice sites are indicated by orange dashed lines, alternative 3′ splice sites by red dotted lines, and alternative transcript end sites by black dash-dot lines. Alternative exons are shown as semi-transparent red rectangles, introns are highlighted in green, and retained introns are marked in dark green. (B) Donut charts showing the distribution of significantly affected alternative splicing events that occur within the ORF, in 6-week vs. 14-month ovaries after PMSG (left) and PMSG + hCG (right). (C) Donut charts showing the distribution of significantly affected alternative splicing events that contain stop codons, in 6-week vs. 14-month ovaries after PMSG (left) and PMSG + hCG (right). (D, E) Horizontal bar plots comparing the location of splicing-induced stop codons in transcripts from affected (blue) and unaffected (orange) ovaries following PMSG (D) or PMSG + hCG (E) treatment. Bars represent the percentage of events resulting in a stop codon in the last exon (left side of each panel) vs. within exons that are not last (right side).

**Figure 5.**
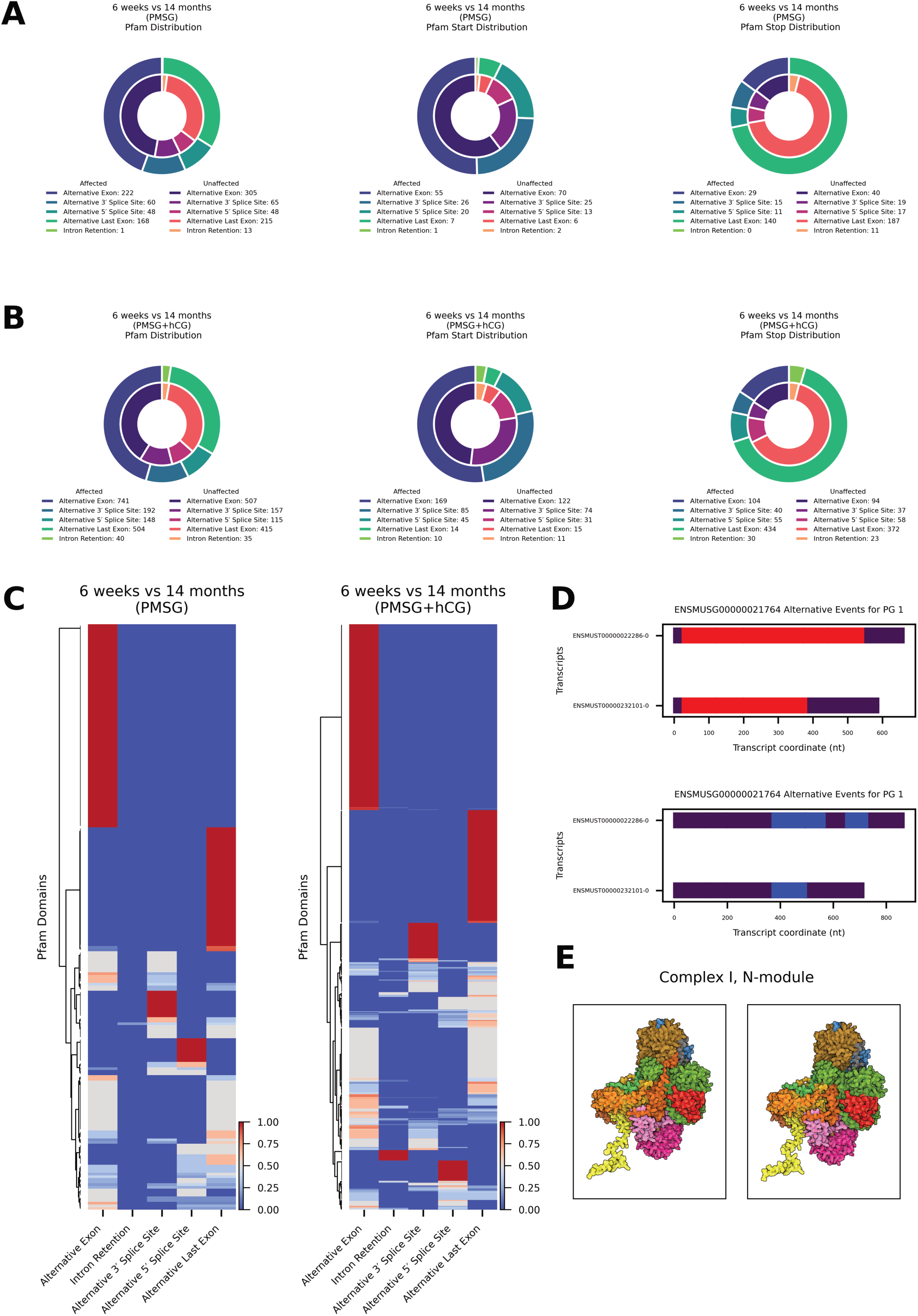
Reproductive aging alters protein domain architecture through splice type-specific remodeling. (A–B) Donut charts showing the distribution of splicing event types associated with Pfam domain gains or losses in significantly affected transcripts of in 6-week vs. 14-month ovaries after PMSG. (A) and PMSG + hCG (B). Each panel depicts the proportion of events contributing to overall domain distribution (left), domain start sites (middle), or domain stop sites (right), categorized by alternative exon, intron retention, alternative 5′ or 3′ splice sites, or alternative last exons. (C) Heatmaps showing the relative enrichment of Pfam domains across splicing event categories in significantly affected transcripts from 6-week vs. 14-month ovaries; left: PMSG, right: PMSG + hCG. Rows represent Pfam domains; columns represent splice event categories. Color scale indicates the normalized fraction of events in which each domain was detected. (D) Transcript architecture of *Ndufs4* gene identified as significantly differentially spliced after PMSG + hCG treatment. Red bars denote the predicted open reading frames. The Pfam domain (blue) is present in the long isoform but absent in the short isoform due to exon skipping. (E) AlphaFold-predicted structures of the mitochondrial Complex I N-module incorporating either the long (left structure) or short (right structure) NDUFS4 isoform. Subunits are color-coded: NDUFS4 is shown in dark orange. The short isoform model lacks the C-terminal portion of NDUFS4, while other subunits retain similar spatial arrangement across both models.

To ensure that shifts in splicing patterns truly reflect biological effects rather than sampling artifacts, we always include both the “affected” and “unaffected” gene sets. The unaffected genes serve as an internal baseline, establishing the background frequency of each alternative splicing event type. By comparing affected vs. unaffected distributions, we can compute valid odds ratios, χ2 statistics (and their standardized residuals), and confidence intervals for each event, thereby identifying which splice classes are genuinely enriched or depleted. Without the unaffected set, any observed deviations could simply mirror overall gene-level biases, undermining the rigor and interpretability of our splicing analysis.

### Data and Code Availability

All command scripts, exact software versions, and detailed runtime parameters are available in the public repository: https://github.com/DrNAlvarez/Dysregulation-of-alternative-splicing-patterns-in-the-ovaries-of-reproductively-aged-mice-. FASTQ files were deposited in the Sequence Read Archive (Leinonen *et al*., 2011), under SRA project ID: PRJNA1263509.

## Results

### Direct RNA-sequencing reveals novel transcripts in the mouse ovary

Direct RNA-sequencing was performed on mRNA from ovaries of mice at two ages (6 weeks-old and 14 months-old), following gonadotropin stimulation with PMSG alone or PMSG followed by hCG (see Methods for details). Across twelve sequencing runs, an average of 3.63 × 10^5^ RNA molecules were sequenced per run (SEM ±1.05 × 10^5^; Figure 1A). Analysis of read lengths showed that the transcripts spanned a wide size range (from a few hundred to over 10^4^ nucleotides), with a mean RNA length of 964 nt (SEM ±0.36 nucleotides; Figure 1B). Notably, the length distribution was highly similar across all experimental conditions (young vs. aged ovaries, with or without hormonal stimulation), indicating a consistent transcript size profile in the ovarian transcriptome.

To comprehensively characterize the ovarian transcriptome, we pooled all direct RNA-sequencing data from the 12 runs and annotated the unique transcripts by Ensembl transcript biotype (Figure 1C, left). In total, 56,862 unique transcripts were identified. A striking majority of these transcripts (72.7%) do not correspond to annotated Ensembl transcripts and were classified as unannotated. Among the annotated fraction, protein-coding transcripts accounted for 19.4%, and retained intron transcripts made up 2.8%. Smaller proportions of known transcript types were detected as well, including long noncoding RNA (lncRNA, 1.3%) and protein-coding transcripts with undefined coding sequence (CDS not defined, 1.3%). All remaining minor categories (each contributing <1% of transcripts) were combined into a single “Other” category, together representing 2.5% of the transcriptome. Because such a large fraction of ovarian transcripts lacked annotation, we next evaluated their protein-coding potential using the Bioseq2seq deep-learning model for mRNA classification (Valencia and Hendrix, 2023). This analysis revealed that 51.6% of the detected transcripts exhibit protein-coding potential, whereas 48.4% are predicted to be noncoding (Figure 1C, right).

### Transcriptome-wide detection of alternative splicing events in the reproductive-aged ovary

An advantage of using direct RNA-sequencing is the ability to assess splicing events at single molecule resolution. We interrogated transcriptome-wide alternative splicing patterns in the mouse ovary by identifying and quantifying splicing events, followed by statistical analysis to determine the significance of observed changes. An example is provided for the gene *Tmem176b* (ENSMUSG000000298) (Figure 2A).

We analyzed alternative splicing within the context of an individual promoter within a gene (see Methods). Our analysis assumes a model of post-transcriptional gene regulation where pre-mRNA generated from a promoter is alternatively spliced into mRNA independently and uniquely, compared to other pre-mRNA generated within the same gene. In Figure 2A, alternative 5’ splice sites are marked with orange dashed lines, alternative 3’ splice sites with red dotted lines, and alternative ends with black dash-dot lines. Alternative exons are highlighted by red shaded rectangles with low opacity, introns are highlighted in green, and highlights in a dark green mark retained introns (not shown). These visual markers help to identify and differentiate between various splicing events and their locations within a promoter group of a gene; Figure 2A is an example of this.

To quantify how reproductive aging affects specific alternative splicing events, for each transcript, we computed the log₂ fold-change (6-weeks vs 14-month PMSG or PMSG + hCG) and statistical significance for a percent spliced (*PS*, see Methods) metric, and propagated that value to each alternative splicing event embedded within the transcript (see Methods). This allowed us to examine event-level changes in inclusion (the transcript contains the event) and exclusion (the transcript skips the event). Panels B–E of Figure 2 summarize *PS* shifts for alternative exons, alternative 5′ and 3′ splice sites, and intron retention.

Following PMSG stimulation (Figure 2B), alternative exon inclusion events showed a strong downward bias: 287 events were down-regulated compared to only 51 up-regulated (binomial *p* < 10⁻^41^), with 509 unaffected. This trend intensified after hCG (Figure 2C), with 760 alternative exon inclusion events down-regulated and 549 up-regulated (*p* = 3 × 10⁻^9^). Alternative 5′ splice site usage was similarly down-biased in both conditions (Figure 2B: 46 down / 13 up, *p* = 2 × 10^-5^; 2C: 104 down / 69 up, *p* = 0.009). Alternative 3′ splice sites inclusion events showed no significant imbalance following PMSG (Figure 2B, *p* = 0.30) but became slightly up-regulated after hCG (Figure 2C: 98 up / 71 down, *p* = 0.045). In contrast, intron retention inclusion events displayed a significant upward shift after hCG (86 up / 26 down, *p* = 1 × 10⁻^8^) but not after PMSG alone (Figure 2B, *p* = 0.29).

For transcripts that excluded annotated events, we observed different trends. Intron retention exclusion events were significantly down-regulated in aged ovaries in both conditions (Figure 2D: 194 down / 78 up, *p* = 1 × 10^-12^; 2E: 418 down / 297 up, *p* = 1 × 10⁻^6^). Alternative exon exclusion events showed no directional bias after PMSG alone (Figure 2D: 61 down / 57 up, *p* = 0.78), but a significant downward shift after hCG (Figure 2E: 297 up / 418 down, *p* = 6 × 10⁻^6^). For alternative 5′ and 3′ splice site inclusion events, aged ovaries showed strong downregulation after PMSG (Figure 2D: alternative 5′ 75 down / 34 up, *p* = 0.0001; alternative 3′ 458 down / 185 up, *p* = 1 × 10⁻^27^), while we observed the opposite following hCG stimulation (Figure 2E: alternative 5′ 294 up / 214 down, *p* = 0.0004; alternative 3′ 1866 up / 1312 down, *p* = 8 × 10^-23^).

We quantified boundary shifts in significantly affected transcripts by comparing proximal vs. distal transcription start sites (TSS) and end sites (TES) in (Fig. 2F,G). Following PMSG stimulation, transcripts whose percent-spliced (*PS*) fold change decreased showed greater proximal TSS usage (OR = 1.50, Fisher’s exact *p* = 0.0004), whereas those with increased *PS* fold change favored distal TESs (OR = 0.29, *p* = 1 × 10^-6^). After hCG stimulation, transcripts with either increased or decreased *PS* fold change were more likely to initiate from proximal TSSs (*PS* > 0: OR = 1.40, *p* = 2 × 10^-5^; *PS* < 0: OR = 1.60, *p* = 1 × 10^-13^), and *PS*-increased transcripts additionally exhibited enhanced distal TES selection (OR = 0.33, *p* = 1 × 10^-27^).

### Reproductive aging globally alters transcriptome complexity

To assess how reproductive aging reshapes transcript complexity and splicing event usage in mouse ovaries following gonadotropin stimulation, we compared the prevalence of promoters that generate single vs. multiple transcripts and the balance of inclusion vs. exclusion events among age-biased (“affected”) and age-neutral (“unaffected”) loci following PMSG or PMSG + hCG treatment (Figure 3).

In PMSG-stimulated ovaries (Figure 3A), aging did not significantly alter promoter-level transcript complexity: aged ovaries yielded 650 single- and 411 multi-transcript promoters at age-biased loci vs. 1104 single- and 662 multi-transcript promoters in age-neutral loci (*χ*^2^ = 0.389, df = 1, *p* = 0.533; Fisher’s OR = 0.948, *p* = 0.522). By contrast, after combined PMSG + hCG (Figure 3B), we observed a marked shift toward multi-transcript promoters, with 1305 multi-vs. 2237 single-transcript promoters at age-biased loci compared to 1405 multi- and 3196 single-transcript promoters in age-neutral loci (*χ*^2^ = 35.57, df = 1, *p* = 2.47 × 10⁻⁹; OR = 0.754, *p* = 2.57 × 10⁻⁹), equivalent to a 1.33-fold increase in the odds of multi-transcript usage with age.

We then asked how aging shifts the global balance of inclusion vs. exclusion across six splicing categories. In PMSG-stimulated ovaries (Figure 3C), aged-affected loci were biased toward inclusion over exclusion at alternative 3′ splice sites (OR = 1.68, *p* = 0.020) and alternative 5′ splice sites (OR = 1.77, *p* = 0.015), while alternative first exons (OR = 0.65, *p* = 5.79 × 10⁻⁸) and intron retention events (OR = 0.22, *p* = 6.26 × 10⁻⁵) was strongly depleted; inclusion vs. exclusion for alternative exons (OR = 0.92, *p* = 0.576) and alternative last exons (OR = 1.20, *p* = 0.113) did not change. Following PMSG + hCG (Figure 3D), inclusion of alternative 5′ ends (OR = 1.36, *p* = 0.024) and alternative last exons (OR = 1.24, *p* = 8.73 × 10⁻⁴) was enriched, whereas alternative first exon inclusion was profoundly depleted (OR = 0.58, *p* = 2.63 × 10⁻³⁷); alternative 3′ splice site (OR = 1.22, *p* = 0.097), alternative exon (OR = 0.96, *p* = 0.528), and intron retention (OR = 1.16, *p* = 0.388) inclusion patterns remained unchanged.

### Reproductive aging influences global stop codon and ORF distribution via alternative splicing

Alternative splicing not only reshapes exon composition, but also rewrites coding potential by modifying open reading frames (ORFs). Figure 4A illustrates these effects for the representative gene *Eef1d* (ENSMUSG00000055762). We quantified how age-biased alternative splicing disrupts ORFs and whether the global pattern of alternative splicing differs from age-neutral loci. Following PMSG stimulation, we detected 759 age-affected events that altered ORFs (Figure 4B, left). Their class distribution diverged significantly from age-neutral events (*χ*^2^ = 15.2, df = 4, *p* = 0.004). Enrichment analysis showed a modest excess of alternative 5′-splice-site usage (OR = 1.39, Fisher’s exact *p* = 0.046) together with a marked depletion of intron-retention events (OR = 0.17, *p* = 6.9 × 10^-4^). Adding hCG increased the number of age-affected ORF-altering events to 2844 (Figure 4B, right). The overall event-class distribution was no longer significantly different from age-neutral genes (*χ*^2^ = 6.64, df = 4, *p* = 0.156), although alternative last-exon usage was modestly under-represented (OR = 0.82, *p* = 7.4 × 10^-4^). Following PMSG stimulation, 245 age-affected and 393 age-neutral stop-codon events were detected (Figure 4C, left). Alternative last-exon usage predominated in both groups (affected 80 %, neutral 75.6 %). The overall class distribution did not differ significantly (*χ*^2^ = 8.48, df = 4, *p* = 0.075). Intron-retention-mediated stops were markedly depleted among age-affected transcripts (2 of 245, 0.8%) compared with age-neutral genes (18 of 393, 4.6 %; OR = 0.17, Fisher’s exact *p* = 8.5 × 10^-3^). With the addition of hCG (PMSG + hCG), 982 age-affected and 983 age-neutral stop-codon events were identified (Figure 4C, right). Class frequencies again showed no significant global difference (*χ*^2^ = 7.39, df = 4, *p* = 0.116). No individual splice-event class exceeded the significance threshold (|standardised residual| < 2), although intron retention displayed a modest, non-significant enrichment in age-affected transcripts (OR = 1.50, *p* = 0.075).

We analyzed the position of premature stop codons following gonadotropin stimulation asking whether termination falls in the annotated last exon (“Last”) or upstream (“Not Last”) for age-affected versus age-neutral transcripts (Figure 4D, E). Following PMSG stimulation (Figure 4D), 223 of 245 stop codons occurred in the last exon of age-affected transcripts; in unaffected transcripts 360 of 393 stop codons occurred in the last exon (*χ*^2^ = 0.012, df = 1, *p* = 0.91). When we examined each splice-event class individually, the global distribution remained non-significant (*χ*^2^ = 8.48, df = 4, *p* = 0.075). Fisher’s exact tests likewise showed no class-specific enrichment of stop codon position among age-affected transcripts: alternative exon (OR = 1.18, *p* = 1.00); alternative 3′ splice site (OR = 0.70, *p* = 0.75); alternative last exon (OR = 0.66, *p* = 0.72); alternative 5′ splice site (OR = 0.38, *p* = 0.36); intron retention (OR = 0, *p* = 0.19). Following PMSG + hCG stimulation (Figure 4E) when stratified by splice-event class, no individual category reached statistical significance after multiple testing correction: alternative exon (OR = 0.52, *p* = 0.057); intron retention (OR = 0.43, *p* = 0.076); alternative 5′-splice-site (OR = 0.59, *p* = 0.24); alternative 3′-splice-site (OR = 0.50, *p* = 0.14); alternative last exon (OR = 1.13, *p* = 0.82). However, globally we observed that in age-affected transcripts 860 stop codons occured in the last exon (87.6 %) and 122 occurred upstream (12.4 %). In contrast, in age-neutral transcripts, 898 of 983 stop codons occurred in the last exon (91.4 %), with 85 located upstream (8.6 %). This modest but significant shift toward upstream stop codon location in age-affected transcripts was confirmed by a 2 × 2 test (*χ*^2^ = 7.04, df = 1, *p* = 0.008; Fisher’s exact OR = 0.67, *p* = 0.0066).

### Reproductive aging alters protein domain architecture through splice type-specific remodeling

To investigate how reproductive aging reshapes protein architecture, we analyzed Pfam domain distribution across alternative splicing across alternative splicing events, following gonadotropin stimulation. Among transcripts with age-affected splicing after PMSG stimulation we detected 499 age-affected alternative splicing events with Pfam domains (Figure 5A, left). The class distribution within the age-affected set was: alternative exon (222; 44.5%), alternative last-exon (168; 33.7%), alternative 3′-splice-site (60; 12.0%), alternative 5′-splice-site (48; 9.6%), and intron retention (1; 0.2%). In age-neutral transcripts we detected 646 Pfam domains; the corresponding profile was: alternative exon (305; 47.3%), alternative last-exon (215; 33.3%), alternative 3′-splice-site (65; 10.1%), alternative 5′-splice-site (48; 7.4%), and intron retention (13; 2.0%). Class composition differed significantly between age-affected and age-neutral sets (*χ*^2^ = 10.63, df = 4, p = 0.031). Fisher’s exact tests showed a significant under-representation of intron-retention events in the age-affected group (OR = 0.098, p = 0.005); no other class was enriched or depleted. For Pfam start-site alterations (Figure 5A, middle) we observed 109 starts in age-affected events and 116 in age-neutral events. Age-affected transcripts were distributed as follows: alternative exon (55; 50.5%), alternative 3′-splice-site (26; 23.9%), alternative 5′-splice-site (20; 18.3%), alternative last-exon (7; 6.4%), intron retention (1; 0.9%). Age-neutral transcripts showed a similar pattern: alternative exon (70; 60.3%), alternative 3′-splice-site (25; 21.6%), alternative 5′-splice-site (13; 11.2%), alternative last-exon (6; 5.2%), intron retention (2; 1.7%). Overall class composition did not differ significantly (*χ*^2^ = 3.50, df = 4, p = 0.48), and no individual class was significantly enriched or depleted (p > 0.13). For Pfam stop-site alterations (Figure 5A, right) we recorded 195 stops in age-affected events vs. 274 stops in age-neutral events. The age-affected set was dominated by alternative last-exon usage (140; 71.8%), followed by alternative exon (29; 14.9%), alternative 3′-splice-site (15; 7.7%), alternative 5′-splice-site (11; 5.6%), and intron retention (0; 0%). Age-neutral transcripts displayed a comparable distribution: alternative last-exon (187; 68.2%), alternative exon (40; 14.6%), alternative 3′-splice-site (19; 6.9%), alternative 5′-splice-site (17; 6.2%), intron retention (11; 4.0%). Class composition did not differ significantly (*χ*^2^ = 8.19, df = 4, p = 0.085). Fisher’s tests confirmed a significant depletion of intron-retention events in the age-affected group (OR = 0, p = 0.0034); all other classes showed no significant differences (p > 0.41).

Following PMSG + hCG stimulation, we detected 1625 age-affected alternative splicing events with Pfam domains (Figure 5B, left). Their class distribution was: alternative exon (741; 45.6 %), alternative last-exon (504; 31.0 %), alternative 3′-splice-site (192; 11.8 %), alternative 5′-splice-site (148; 9.1 %), and intron retention (40; 2.5 %). In the 1229 age-neutral Pfam domains, the corresponding profile was: alternative exon (507; 41.3 %), alternative last-exon (415; 33.8 %), alternative 3′-splice-site (157; 12.8 %), alternative 5′-splice-site (115; 9.4 %), and intron retention (35; 2.8 %). Overall class composition did not differ significantly between age-affected and age-neutral sets (*χ*^2^ = 5.64, df = 4, *p* = 0.23). However, Fisher’s exact test indicated a modest enrichment of alternative exon events in the age-affected group (OR = 1.19, *p* = 0.022), whereas the other classes showed no significant differences (*p* > 0.12). For Pfam start-site alterations detected after PMSG + hCG stimulation (6-weeks vs 14-months; Figure 5B, middle), we observed 323 starts in age-affected events and 253 starts in age-neutral events. Within the age-affected set the distribution was: alternative exon (169; 52.3 %), alternative 3′-splice-site (85; 26.3 %), alternative 5′-splice-site (45; 13.9 %), alternative last-exon (14; 4.3 %), and intron retention (10; 3.1 %). The age-neutral transcripts showed a comparable profile: alternative exon (122; 48.2 %), alternative 3′-splice-site (74; 29.2 %), alternative 5′-splice-site (31; 12.3 %), alternative last-exon (15; 5.9 %), and intron retention (11; 4.3 %). Overall class composition did not differ significantly between age-affected and age-neutral groups (*χ*^2^ = 2.54, df = 4, *p* = 0.64), and Fisher’s exact tests revealed no significant enrichment or depletion for any individual class (*p* > 0.35 for all comparisons). For Pfam stop-site alterations detected after PMSG + hCG stimulation (Figure 5B, right) we identified 663 stops in age-affected events and 584 age-neutral events. In the age-affected set the distribution was: alternative last-exon (434; 65.5%), alternative exon (104; 15.7 %), alternative 5′-splice-site (55; 8.3 %), alternative 3′-splice-site (40; 6.0 %), and intron retention (30; 4.5 %). Age-neutral events showed a comparable profile: alternative last-exon (372; 63.7 %), alternative exon (94; 16.1 %), alternative 5′-splice-site (58; 9.9 %), alternative 3′-splice-site (37; 6.3%), and intron retention (23; 3.9 %). Overall class composition did not differ significantly between age-affected and age-neutral groups (*χ*^2^ = 1.40, df = 4, *p* = 0.84), and Fisher’s exact tests confirmed that no individual event class was significantly enriched or depleted (*p* > 0.32 for all comparisons).

Hierarchical clustering of Pfam domains segregate into distinct splice-event–specific blocks (Figure 5C). Following PMSG stimulation (left heat-map) 222 unique Pfam domains were identified in age-affected transcripts. Most domains showed their strongest perturbation through alternative exon events (126; 56.8 %), forming a large dominant cluster. A second block consisted of domains preferentially altered by alternative last exon usage (57; 25.7 %), while smaller branches were driven by alternative 3′ splice site (21; 9.5 %), alternative 5′ splice site (18; 8.1 %) and intron retention (1; 0.4%). After combined PMSG + hCG stimulation (right heat-map) the landscape expanded to 608 unique Pfam domains within age-affected transcripts. The overall architecture was preserved: domains still partitioned mainly into alternative exons (328; 53.9 %) and alternative last exon (160; 26.3 %) clusters, with additional branches for alternative 3′ splice site (62; 10.2 %) and alternative 3′ splice site (43; 7.1 %) events. A new, smaller module emerged in which (15; 2.5 %) were preferentially perturbed by intron retention.

Hierarchical clustering of the Pfam-by-splice-event matrices (Figure 5C) showed that several of the age-segregating clusters were dominated by domains involved in mitochondrial function, including multiple Pfam domains contributing to Complex I function (NDUS4, NDUFA5, NDUFB2, NDUFB8, NADHdh_A3), assembly factor domains (UQCC3, TMEM70), cristae-organising MICOS component domains (Mitofilin, MIC19/MIC25), inner-membrane carrier domains (Mito_carr), and TIM/TOM import machinery domains (Tim17, Tom5). The presence of the NDUS4 domain in one of the strongest clusters dovetails with our splice-isoform analysis of *Ndufs4* itself. Following PMSG + hCG stimulation, *Ndufs4* is differentially spliced during aging, showing ≈ 32-fold and ≈ 4-fold increases in a long and short isoform respectively (Figure 5D). The longer isoform retains the full NDUS4 Pfam domain, while the shorter isoform lacks this region. AlphaFold 3 modelling of the mitochondrial Complex I N-module (Figure 5E) indicates that incorporation of the truncated NDUFS4 short isoform yields a visibly shortened subunit (dark orange) without major rearrangement of the surrounding proteins.

## Discussion

### Alternative splicing landscape in aged ovaries following gonadotropin stimulation

Our long-read transcriptome analysis reveals that reproductive aging is accompanied by extensive alterations in mRNA splicing and isoform usage in the ovary. When comparing young vs. aged mice subjected to gonadotropin stimulation (PMSG with or without ovulatory hCG), we observed a broad remodeling of transcript isoforms. Aged ovaries expressed a markedly expanded isoform repertoire per gene, indicating greater transcriptomic heterogeneity with age. In practical terms, aged ovaries produce more alternative mRNA variants than their young counterparts, reflecting pervasive changes in splice site selection and transcript start/end usage. We detected widespread age-biased splicing events, including exon skipping, alternative donor/acceptor site usage, and intron retention, that differentiated 14-month ovaries from 6-week ovaries (Figures 2, 3). Notably, these shifts were stimulus-dependent: after PMSG alone, hundreds of alternative exon inclusion events were already significantly skewed in aged ovaries (predominantly decreased inclusion, i.e. more exon skipping in the aged group), but the differences became even more pronounced after the full PMSG + hCG treatment. The LH-like surge (hCG) unveiled more extreme splicing biases: aged ovaries exhibited a greater suppression of exon inclusion and a significant increase in aberrant intron retention events relative to young ovaries. This suggests that the acute hormonal and transcriptional demands of ovulation exacerbate underlying splicing vulnerabilities in the aged ovary. In addition, aging was associated with shifts in transcript boundary selection; aged ovaries favored more distal transcription start sites and alternative 3′ ends in many transcripts (Figure 2F, G), which could modify 5′/3′ UTR lengths and thereby influence mRNA stability or translation. Together, these findings from Figures 1-5 paint a picture of an aging ovary undergoing not just transcriptional changes in magnitude, but qualitative changes in isoform output for numerous genes.

Crucially, many age-related splicing changes are predicted to have functional consequences at the protein level. We found that a substantial fraction of altered splicing events in aged ovaries changed the mRNA open reading frame. In many transcripts, the inclusion or exclusion of specific exons in older animals introduced premature stop codons or frame shifts, or caused the loss of important protein-coding segments (Figures 4A-E). Consequently, the aged ovary tends to produce a higher proportion of truncated proteins or proteins with altered domain architectures. For instance, several transcripts in aged ovaries utilized alternative last exons or extended 3′ UTRs that added or removed regulatory motifs, and numerous exon-skipping events deleted internal coding sequences. Such isoform switches can disrupt conserved protein domains and interaction motifs. The net result is that the proteome of the aged ovary is qualitatively different, potentially less functional, despite the source genes being the same. This underscores that reproductive aging involves a post-transcriptional dimension: it is not only changes in gene expression levels, but which transcript isoforms are expressed that differs in young vs. aged ovaries. This principle aligns with observations in other systems that aging broadly correlates with dysregulated RNA splicing patterns and isoform diversity across multiple tissues (Guantes *et al*., 2015; Lee *et al*., 2016; Wang *et al*., 2018). Our data extend this paradigm to ovarian aging, highlighting that the ovary follows the general rule that advanced age brings about pervasive alternative splicing shifts in genes linked to fundamental aging processes.

### Isoform diversity, mitochondrial function, and the “splicing-energy-aging” axis

One of the most intriguing patterns emerging from our dataset is the presence of metabolic and mitochondrial genes among those with age-dependent splicing alterations (Figure 5). Transcripts encoding subunits of the mitochondrial electron transport chain (ETC) and other metabolic enzymes were disproportionately affected by alternative splicing in aged ovaries. This observation suggests a mechanistic link between ovarian energy metabolism and splicing regulation in the context of aging. Indeed, there is growing evidence for a splicing-energy-aging axis in biology (Ferrucci *et al*., 2022). Ferrucci and colleagues have hypothesized that declining mitochondrial function and cellular bioenergetic stress can actively drive changes in pre-mRNA splicing as an adaptive response, a concept termed the “energy-splicing resilience axis” of aging. In this model, when ATP is plentiful, cells maintain a normal, basal, splicing program, but under conditions of energy deficit (such as in aged tissues with mitochondrial dysfunction), cells broaden their repertoire of spliced isoforms as a potential resilience or compensatory mechanism (Ferrucci *et al*., 2022). Our findings provide an opportunity to examine this hypothesis in the ovarian context. The aged ovary is known to experience metabolic challenges; aged oocytes and granulosa cells have impaired mitochondrial ATP production and higher reactive oxygen species (ROS) levels (Keefe *et al*., 1995; Barritt *et al*., 2000; Chan *et al*., 2005), along with reduced metabolic flexibility (Bao *et al*., 2024). Consistent with the energy-splicing axis idea, we observed that many of the splicing changes in aged ovaries occur in genes related to mitochondrial oxidative phosphorylation, antioxidant defenses, and other stress response pathways, as has been reported previously (Zhang *et al*., 2019). Prior studies across tissues have noted that age-associated alternative splicing frequently targets genes involved in metabolism, mitochondria, and apoptosis (Lee *et al*., 2016), and our ovarian data echo those findings. For example, an aging study in mice reported that exon-skipping events were frequent in genes tied to mitochondrial functions and inflammation (Stilling *et al*., 2014). Furthermore, experimental work in cell culture has shown that inducing mitochondrial dysfunction (and hence ATP depletion) can *directly* trigger splicing shifts: in neuronal cells, chemical inhibition of mitochondrial respiration caused global production of shorter mRNA isoforms, and importantly this splicing modulation correlated with ATP loss rather than with ROS generation alone (Maracchioni *et al*., 2007). Such evidence underscores that the splicing machinery is sensitive to cellular energy status.

Applying this framework to our results, we propose that the aged ovary, faced with chronic energy stress, may activate or succumb to a splicing response akin to other energy-deprived cells. On one hand, increased isoform diversity in aged ovaries could be an attempt at resilience. The production of alternative isoforms, some of which may be tailored to low-energy conditions, may be an adaptive response of ovarian cells to maintain homeostasis. This outcome would align with the concept of splicing as a stress-response mechanism to maintain homeostasis (Ferrucci *et al*., 2022). Thus, the isoform reprogramming in aged ovaries may partly represent an adaptive remodeling intended to cope with mitochondrial insufficiency and oxidative stress that accompany reproductive aging.

However, splicing dysregulation might also contribute to pathology. Our data provide clear examples of potentially maladaptive splice variants emerging in aged ovaries that could impair cellular function. In particular, transcripts encoding critical mitochondrial proteins appear to be vulnerable to aberrant splicing, raising concern that some splicing changes actively undermine the bioenergetic capacity of the ovary. The clearest case we uncovered is that of *Ndufs4*, a gene encoding a small (18 kDa) accessory subunit of mitochondrial Complex I. Complex I (NADH:ubiquinone oxidoreductase) is the first and largest complex of the respiratory chain, and its proper assembly is vital for electron transport and ATP generation. We found that aging caused an increase in the short *Ndufs4* isoform that lacks the canonical C-terminal Pfam domain present in the long isoform (Figure 5D). In aged ovaries, after PMSG + hCG treatment, the short *Ndufs4* transcript was upregulated (∼4-fold) compared to young. Our AlphaFold modeling of Complex I N-module suggests that the short NDUFS4 removes a critical β2–β3 loop that normally passes under NDUFS1 and links NDUFS4 with the subunits NDUFA12, NDUFA6, and NDUFS9, a structural junction essential for stabilizing the NADH-dehydrogenase (N) module (Yin *et al*., 2024). As a result, the N module may only loosely associate, potentially failing to recruit NDUFA12, causing complex I to stall at an immature assembly intermediate and prevent full enzyme maturation, impairing its function (Yin *et al*., 2024). This splice-driven change is likely to have concrete repercussions for mitochondrial function in the aged ovary. It is well established that *Ndufs4* is essential for the stability of Complex I: mice completely lacking NDUFS4 cannot form fully functional Complex I, leading to drastically reduced Complex I activity, severe neurodegeneration, growth stunting, and premature death (Kruse *et al*., 2008). Even partial loss of NDUFS4 has significant effects. In cell culture, *Ndufs4*-null fibroblasts accumulate incomplete Complex I assemblies and exhibit elevated ROS levels due to electron leak from the dysfunctional complex (Valsecchi *et al*., 2013). *In vivo*, *Ndufs4* knockout mice exhibit systemic metabolic deficiencies; notably, female *Ndufs4*^−/−^ mice are infertile and have smaller ovaries, and their oocytes or zygotes show poor developmental competence, underscoring that *Ndufs4* is required for normal ovarian function and early embryonic development (Wang *et al*., 2017). Thus, a switch toward a nonfunctional NDUFS4 isoform in aged ovaries could effectively phenocopy a mild Complex I deficiency. We propose that this splicing change potentially contributes to the mitochondrial dysfunction known to plague the aging ovary (Zhang *et al*., 2019). Ovarian aging is strongly associated with increased oxidative stress in oocytes and granulosa cells (Sasaki *et al*., 2019). Our results provide a novel molecular explanation for this phenomenon: mis-splicing of a Complex I subunit (and potentially other ETC components) can directly elevate ROS and trigger a downward spiral of cellular damage in the ovary.

Interestingly, NDUFS4 may not be unique in this respect; our Pfam domain analysis (Figure 5A–C) indicates that multiple Pfam domains belonging to mitochondrial proteins (especially ETC subunits) are altered via alternative splicing in the aged ovary. In other words, *Ndufs4* is one example of a broader pattern in which the protein domain architecture of mitochondrial complexes is being altered by splicing in aged ovaries. We speculate that such changes, if they result in unstable or less efficient complexes, could cumulatively compromise cellular energy production within the ovary. It is worth noting that alternative splicing of *Ndufs4* is not just a singular observation in our study, but may reflect a regulated mechanism. For example, the scaffold protein IQGAP1 was shown to influence increased skipping of exon 2 in *Ndufs4* pre-mRNA resulting in an NMD inducing isoform, thereby tuning Complex I activity in response to cellular cues (Papadaki *et al*., 2023). In summary, the *Ndufs4* case exemplifies how an age-altered splicing event might be causally linked to a key aging hallmark (mitochondrial ROS accumulation) in the ovary. This is an instance where an alternative isoform is not merely a biomarker of aging, but potentially a driver of functional decline.

### Conclusion

By illuminating the landscape of isoform remodeling in the aging mouse ovary, our study adds a new dimension to our understanding of reproductive aging. It also raises several important questions. First, what are the upstream causes of the splicing dysregulation in aged ovaries? Second, which of the identified splice isoform changes are truly driving functional decline versus which are neutral byproducts of aging? How do these splicing changes manifest under natural reproductive aging conditions? Finally, this work prompts a broader reconsideration of how we frame ovarian aging. Traditionally, discussions of ovarian aging focus on the decline in follicle quantity and the intrinsic defects in aged oocytes (chromosomal aneuploidy, mitochondrial insufficiency, etc.). Our findings add mRNA splicing fidelity and isoform balance as critical factors in the health of the ovarian microenvironment. They suggest that ovarian aging is not a linear, unidirectional deterioration but a dynamic process involving the cell’s gene regulatory network attempting to respond, successfully or not, to mounting stress. In practical terms, this insight opens new avenues: if aberrant splicing is contributing to ovarian aging, could interventions that support the splicing machinery or improve mitochondrial function mitigate some of the age-related decline? For example, enhanced expression of certain splicing factors has been shown to improve cell function in aged somatic tissues (Latorre *et al*., 2017); whether such strategies could extend the reproductive lifespan is an intriguing idea.

In conclusion, our study demonstrates that alternative splicing undergoes systematic reprogramming in the aging ovary, with potentially significant biological and mechanistic consequences. We hypothesize that the observed post-transcriptional remodeling is both a response to the cellular stresses of aging and a potential driver of further dysfunction. By linking specific splice variants (like the *Ndufs4*) to known aging phenotypes (mitochondrial ROS and infertility), we provide a proof-of-concept that isoform changes could be more than molecular noise, but may be pivotal pieces of the aging puzzle. These findings urge a more integrative view of reproductive aging, one that includes the nuanced layer of RNA splicing regulation alongside genetics, epigenetics, and metabolism. Moving forward, dissecting the splicing–energy–aging axis in the ovary will be critical for a deeper understanding of ovarian longevity and could identify novel targets to ameliorate age-related fertility decline. The ovary, as one of the earliest-aging organs, might also serve as a valuable model for studying how cellular resilience mechanisms like alternative splicing unfold with age in other tissues. Ultimately, unraveling how and why splicing patterns shift with age will deepen our understanding of ovarian aging and may suggest new strategies to preserve female reproductive health in the face of advancing age.

## Author contribution statement

Conceptualization, P.B. and N.S.A.; methodology, P.B. and N.S.A.; validation, P.B. and N.S.A.; computational analysis, N.S.A.; investigation, P.B. and N.S.A.; writing—original draft preparation, P.B. and N.S.A.; writing— review and editing, P.B. and N.S.A. All authors have read and agreed to the published version of the manuscript.

## Acknowledgements

Not applicable.

